# A topological analysis of difference topology experiments of condensin with Topoisomerases II

**DOI:** 10.1101/795898

**Authors:** Soojeong Kim, Isabel K. Darcy

## Abstract

An experimental technique called *difference topology* combined with the mathematics of tangle analysis has been used to unveil the structure of DNA bound by the Mu transpososome. However, difference topology experiments can be difficult and time-consuming. We discuss a modification that greatly simplifies this experimental technique. This simple experiment involves using a topoisomerase to trap DNA crossings bound by a protein complex and then running a gel to determine the crossing number of the knotted product(s). We develop the mathematics needed to analyze the results and apply these results to model the topology of DNA bound by 13S condensin and by the condensin MukB.

**SUMMARY STATEMENT:** Tangles are used to model protein-DNA complexes: A 3-dimensional ball represents protein while strings embedded in this ball represent protein-bound DNA. We use this simple model to analyze experimental results.

## INTRODUCTION

Proteins bind DNA in many genetic activities such as replication, transcription, packaging, repair, and rearrangement. Understanding the DNA conformation within protein-DNA complexes is useful for modeling and analyzing reactions (Crisona et al., 1999; Harshey and Jayaram, 2006; Kumar et al., 2017; Kimura et al., 1999; Pathania et al., 2002; Petrushenko et al., 2006). Laboratory techniques have been developed to study the shape of protein-DNA complexes including X-ray crystallography, cryo-EM (cryogenic electron microscopy), AFM (atomic force microscopy) and NMR (nuclear magnetic resonance). Technology has significantly advanced, but it is still unsuccessful for large complexes in which proteins bind multiple DNA segments. To study protein-bound DNA, an experimental technique called *difference topology* combined with the mathematics of tangle analysis has been used (Kimura et al., 1999; Pathania et al., 2002; Petrushenko et al., 2006; Darcy et al., 2009; Kim and Darcy, 2015). Note this technique focuses on determining the topology of DNA bound in a protein complex. Tangle analysis ignores the shape of the protein and cannot determine the exact geometry of the protein-bound DNA. But this simplified model has been used to determine reaction pathways and as a basis for more complex models (Pathania et al., 2003; Vazquez et al., 2005; Harshey and Jayaram, 2006; Grainge et al., 2007; Darcy et al., 2008; Shimokawa et al., 2013; Stolz et al., 2017). However, difference topology experiments can be difficult and time-consuming.

We discuss a modification that greatly simplifies this experimental technique. We develop the mathematics needed to analyze the results. We apply these results to experiments performed on 13S condensins by Kimura et al. (Kimura et al., 1999) and the condensin MukB by Petrushenko et al.(Petrushenko et al., 2006).

Tangles were used to model the biological results in (Pathania et al., 2002). An *n-string tangle* is a three-dimensional ball with *n* strings properly embedded in it. When a protein complex binds DNA at *n* sites, the DNA-protein complex can be modeled by an *n*-string tangle (Ernst and Sumners, 1990). Fig. 1A shows the 3-string tangle model for the Mu transpososome as determined in (Pathania et al., 2002). The protein is modeled by the ball and the protein-bound DNA segments are modeled by the strings. Note that this is a 2-dimensional model. But 2-dimensional tangle models can be used to create a 3-dimensional tangle model (Vazquez et al., 2005).

**Fig. 1.**
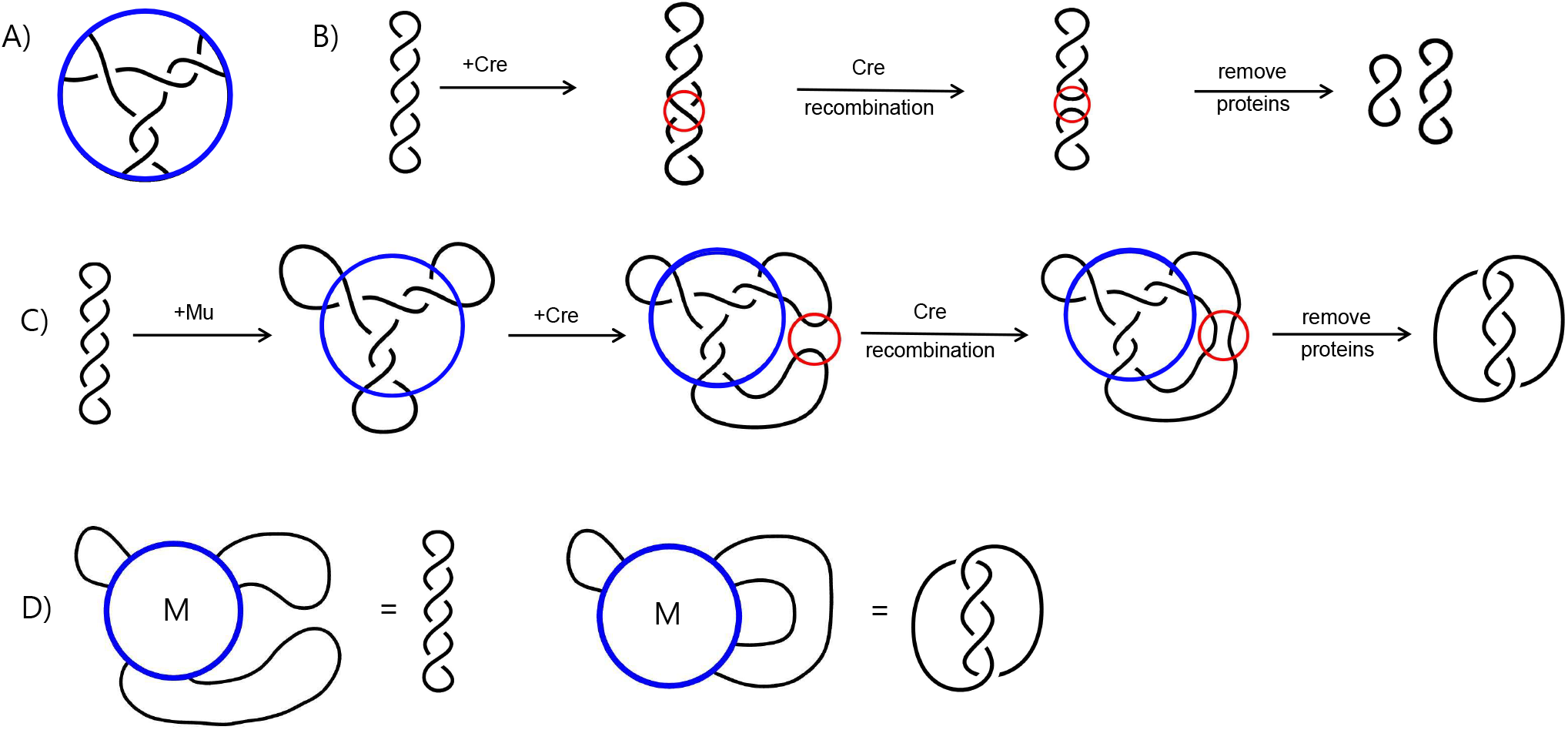
Difference Topology. (A) The topology of DNA bound within the Mu transpososome. Note that Mu is represented by the blue circle and the three black lines represent the three DNA segments bound by Mu. (B) Cre acting on the DNA substrate. Cre is represented by the red circle. Note that the substrate is unknotted negatively supercoiled DNA. DNA is generally underwound and hence forms right-handed negative supercoils. The product of Cre recombination in this case is two smaller unlinked circles. (C) An example of a difference topology experiment from (Pathania et al., 2002). Mu is represented by the blue circle while Cre is represented by the smaller red circle. Note the product of Cre acting on DNA bound by Mu is a 4-crossing catenane, while Cre acting on DNA without Mu produces two small unlinked DNA circles. This difference is the result of crossings bound by Mu that are trapped by Cre recombination. (D) Tangle equation modeling the reaction in (C). The substrate equation is shown on the left and the product equation is shown on the right.

Difference topology experiments can be used to create a system of tangle equations where one can solve for the topology of the DNA bound by protein. The system of two tangle equations modeling the reaction in Fig. 1C is given in Fig. 1D. The unknown variable *M* represents the tangle modeling the Mu transpososome before the tangle solution was determined. Before Cre recombination, the DNA is unknotted. After Cre recombination, the DNA product is a 4-crossing catenane. Note that the tangle in Fig. 1A is one solution for the tangle variable *M* in this system of two tangle equations. However this pair of equations is insufficient to determine a unique tangle solution modeling Mu. Thus Pathania et al performed many additional experiments in order to create a system of nine tangle equations to solve for the unknown tangle *M*. Using only three of these equations, it was proved both mathematically (Darcy et al., 2009) and computationally (Darcy et al., 2006) that the model shown in Fig. 1A is the only biological reasonable solution.

One issue with the difference topology technique using a recombinase such as Cre is the many experiments that need to be performed in order to create an accurate system of tangle equations. The sites for Cre can be placed in two different types of orientations, directly versus inversely repeated. The two different orientations are used to determine the topology of the outside loops. See (Pathania et al., 2002) for more details. Moreover, the location of binding sites for Cre must be determined. If the binding sites for Cre are placed too close to the binding sites for Mu, Cre will be unable to act. If the Cre binding sites are placed too far from the binding sites for Mu, recombination will trap extra DNA crossings, resulting in a variety of different types of DNA knots or catenanes. Thus many experiments were performed in order to obtain unique products in each experiment used to create tangle equations. Next we will discuss how topoisomerase can be used instead to greatly simplify this experimental technique.

## Results

Instead of performing multiple experiments to generate a few tangle equations, one can perform a single experiment to generate multiple tangle equations by using a type II topoisomerase instead of a recombinase. Type II topoisomerase will bind to one segment of DNA, break it, and allow a second segment to pass through this break before resealing the break. Thus they can change DNA topology. Type II topoisomerases are not site-specific; that is, they can bind anywhere along DNA. Using examples, we illustrate some modeling challenges and discuss benefits of using a topoisomerase instead of a recombinase to trap crossings when performing difference topology experiments.

### Modeling two DNA segments bound by protein via difference topology using topoisomerase

Suppose we wish to study a protein that binds two segments of DNA as shown in Fig. 2 where the protein under study is represented by the tangle *P*. Protein complexes that bind more than two DNA segments will be discussed in later sections. As before, we do both the control experiment where topoisomerase acts on naked DNA (Fig. 2A) as well as one where protein *P* is added first so that it binds DNA before topoisomerase is added. (Fig. 2B). Note that the control experiment is very important to confirm that reaction conditions are such that topoisomerase action does not result in knotted DNA unless protein *P* has been added first. Topoisomerase under normal circumstances will unknot knotted DNA (Leroy et al., 1980), but under some circumstances (such as in a difference topology experiment, but also under some reaction conditions (Wasserman and Cozzarelli, 1991)), topoisomerase will knot DNA. So the control experiment must be done to ensure that the knots produced by topoisomerase in a difference topology experiment are due to the presence of protein *P* and not topoisomerase acting on naked DNA.

**Fig. 2.**
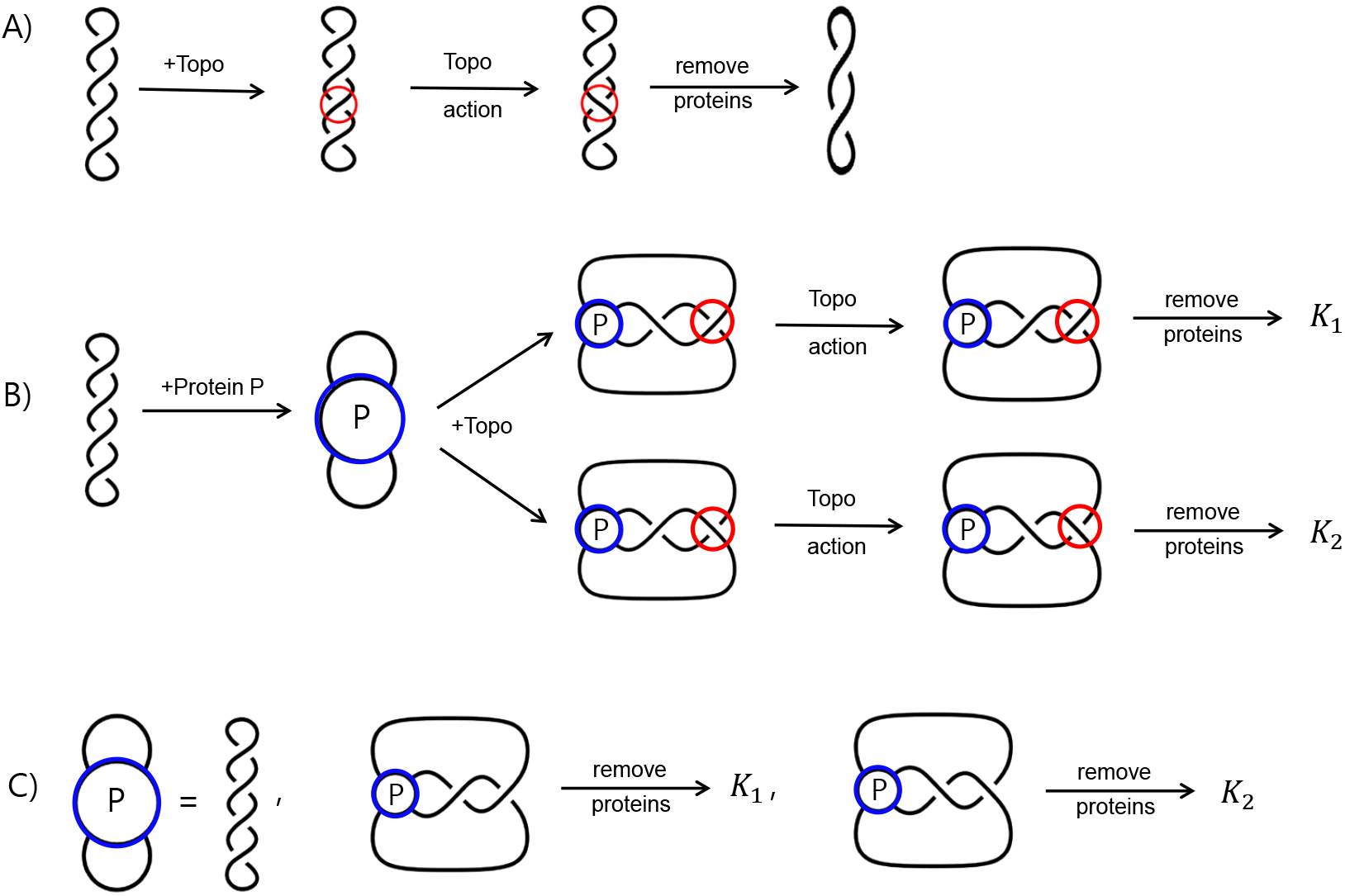
Difference Topology using a topoisomerase. (A) Topoisomerase acting on unknotted supercoiled DNA. Topoisomerase is represented by the red circle. Reaction conditions are chosen so that topoisomerase does not create DNA knots. (B) An example of a difference topology experiment using a topoisomerase where *K*_1_ and *K*_2_ refer to the knotted products. The protein under study is modeled by the blue circle labeled *P* while topoisomerase is represented by the red circle. (C) Tangle equations modeling the reactions in (B). The first equation corresponds to the unknotted substrate equation. The middle equation is the product equation where topoisomerase action is represented by a left-handed clasp, while the last equation is the product equation where topoisomerase action is represented by a right-handed clasp.

In Fig. 2B, the protein *P* is added first, binding DNA segments. Topoisomerase is then added to the reaction. The two loops come together forming two DNA crossings. Note that there are two different ways that the loops can come together as shown in this figure, resulting in two cases. Topoisomerase changes one of the two crossings. Without loss of generality, mathematically speaking, topoisomerase is shown acting on the crossing on the right in both cases. Three possible tangle equations modeling the reaction in Fig. 2B are shown in Fig. 2C. Note that we would still obtain these same tangle equations if topoisomerase acted on the crossing on the left in both cases. In the middle figure of Fig. 2C, topoisomerase action is modeled by a left-handed clasp, while the figure to the right shows a right-handed clasp. Whether or not topoisomerase action is modeled by a left-handed or right-handed clasp can be projection dependent as shown in Movie 2. For example, the two segments bound by topoisomerase could cross at a 90° angle. After topoisomerase action, if one views the 3-dimensional model from the right, one would see a left-handed clasp; while if one viewed it from the left, one would see a right-handed clasp. Note that choosing a projection fixes the handedness of the clasp. Which projection we take also affects the 2-dimensional tangle solutions for *P* (Vazquez et al., 2005). Thus once we choose a particular projection to represent topoisomerase action (i.e., clasp handedness), we have also chosen a projection for the 2-dimensional tangle *P*.

Whether we should use all three tangle equations shown in Fig. 2C depends on the 3-dimensional conformation of the DNA loops emanating from the protein complex. Consider first the unknotted substrate equation (the first equation in Fig. 2C). In this equation showing the unknotted substrate, none of the three loops interact. This is the same assumption that was made by Pathania et al (Pathania et al., 2002) when using Cre recombinase to determine the DNA conformation within the Mu transpososome. This is a reasonable assumption as illustrated in Movie 1. Depending on how the 3-dimensional protein-DNA complex is projected, we may or may not see two of the loops cross. In most cases, there should be a projection where the loops do not cross as assumed in Fig. 2C. However, if it is later determined that no such projection exists, the solution found using the outside loop configuration shown in Fig. 2C can be easily modified to satisfy a different configuration of outside loops (Darcy et al., 2006; 2009). Thus we will always assume that when protein *P* binds DNA, the loops emanating outside of the protein-DNA complex do not cross in the substrate equation (per Figs 1D, 2C, and Eqn 1 in Table 1).

**Table 1.**
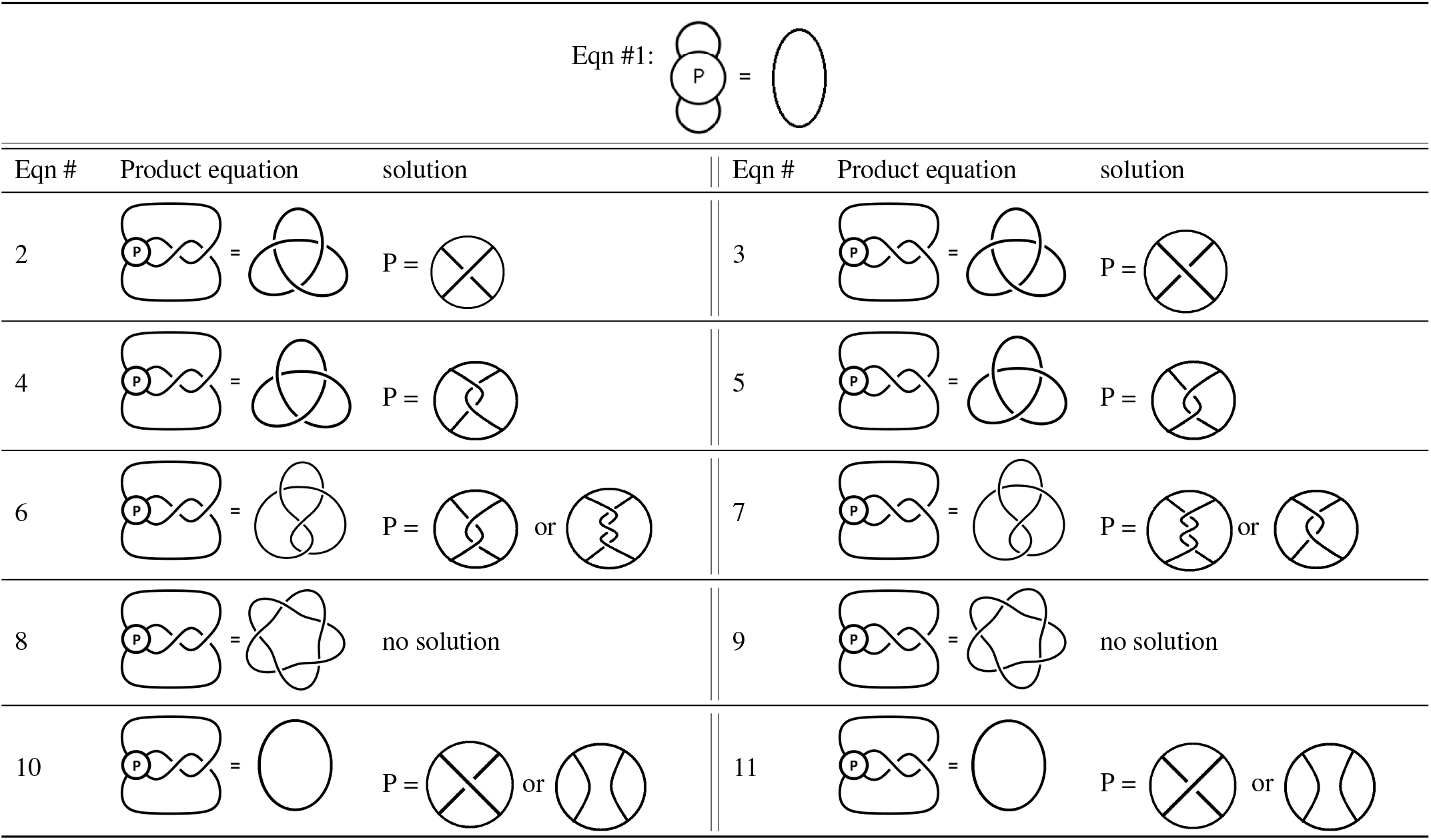
The first row shows Eqn 1, the unknotted substrate equation. For the remaining rows, the first and fourth columns give the equation number. The second and fifth columns show possible tangle equations modeling potential knotted products. The third and sixth columns show solutions to the system of two tangle equations involving the substrate equation (Eqn 1) and one of the product equations.

The main question is whether we should use both of the product equations. In one case, topoisomerase creates a right-handed clasp. In the other case, topoisomerase creates a left-handed clasp. How the loops emanate from the protein-DNA complex could make one configuration more likely than the other one. Thus one must allow for some ambiguity in terms of which tangle equations best model the difference topology experiment. However, we can still gain useful information despite this ambiguity.

Table 1 shows solutions to systems of two tangle equations where the first equation is always the unknotted substrate equation shown at the top of this table and the second equation is one of the product equations. One can also use the TopoICE-X (Darcy et al., 2008) and TopoICE-R (Darcy and Scharein, 2006) software within KnotPlot (Scharein, 1998) to solve most biologically relevant 2-string tangle equations. We will illustrate how Table 1 can be used with the examples below. In these examples, we will sometimes assume we only know the crossing number of the knotted product as this can be determined via gel electrophoresis. The crossing number of a knot is the smallest number of crossings needed to draw that knot. The unknot is the only knot that can be drawn with fewer than three crossings. There are two three crossing knots and one four crossing knot (shown in Table 1). For a table that includes higher crossing knots, please see (Rolfsen, 1976) and https://knotplot.com/zoo/.

#### Example 1

Suppose that the products observed are 3-crossing and 4-crossing knots. Thus assuming protein *P* binds a unique DNA conformation, we have a system of three tangle equations (per Fig. 2C): one for the unknotted substrate and one each for the two products. From Table 1 these equations must be Eqns 1, 5, 6 or Eqns 1, 4, 7. If we look at any other combination of equations, we do not have a common solution. For example, Eqns 1 and 2 mean protein *P* must bind one crossing. But if *P* binds only one crossing, topoisomerase action cannot produce a 4-crossing knot per Eqns 6 and 7. Per these equations, protein *P* must bind to two or three crossings in order to produce a 4-crossing knot when starting with an unknotted substrate. Thus Eqn 2 cannot be one of the equations modeling this reaction if protein *P* binds a unique DNA conformation. However, the system of three tangle Eqns 1, 5, and 6 has a solution where protein *P* binds two negative supercoils. Note that this two crossing solution satisfies the system of two tangle Eqns 1 and 5 as well as the pair of tangle Eqns 1 and 6 and thus satisfies all three Eqns, 1, 5, and 6. If Eqns 1, 4, and 7 model these reactions, then protein *P* binds to two positive supercoils.

One should also consider the possibility that one of these products is the result of multiple rounds of topoisomerase action. For example, topoisomerase could act twice on the unknotted substrate to produce the 4-crossing knot. However it is not possible to convert a 3-crossing knot to a 4-crossing knot (or vice versa) via a single topoisomerase action (Darcy and Sumners, 1997; Torisu, 1998; Darcy et al., 2008). Since no other knot types were detected, it is unlikely that topoisomerase acted more than one time on the unknotted substrate to produce the 4-crossing knot (and similarly for the 3-crossing knot).

#### Example 2

Suppose that the only products are 3-crossing knots. The tangle equations modeling this reaction would be the substrate equation (Eqn 1 in this table) along with only one of the Eqns 2-5. Any system of three equations involving Eqn 1 and two equations from Eqns 2-5 has no solution as these equations do not have a solution in common. Thus only one of equations 2 - 5 can hold. Hence we can only determine that protein *P* binds to one or two crossings. If we wish to determine the handedness of these crossings, one would need to identify whether the 3-crossing knots are right-handed or left-handed via AFM or EM. But since only one of these equations can hold, that means that topoisomerase action in this case must result in either the left-handed or right-handed clasp, but not both. Hence we can infer that the loops must emanate from protein *P* in such a manner that only one-handedness is possible.

#### Example 3

Suppose one of the products is the five-crossing knot shown in Eqns 8 and 9. Topoisomerase must act twice on unknotted DNA to produce this five-crossing knot. Thus there is no solution to the system of two tangle Eqns 1 and 8 and well as Eqns 1 and 9. We can often distinguish between products which require two or more rounds of topoisomerase action from those which can be produced via a single crossing change via topoisomerase action (Darcy and Sumners, 1997; Torisu, 1998; Darcy et al., 2008). However, knotted products that require multiple rounds of topoisomerase action can still be used for modeling. We will discuss this example further when we analyze experiments involving condensins.

#### Example 4

Suppose the only knot types detected are unknots. Then it is possible that the shape of DNA bound by protein is simple per Eqns 1 and 10/11 where the protein complex binds at most one crossing. However, it is also possible that the protein complex under study did not stably bind DNA and thus the reaction shown in Fig. 2B may not have occurred. Thus if no knots are detected, one cannot make any conclusions regarding the DNA conformation bound by protein.

### Modeling three DNA segments bound by protein via difference topology using topoisomerase

The DNA within the Mu transpososome is modeled by a 3-branched supercoiled structure as shown in Fig. 1A. A more general 3-branched structure is shown in Fig. 3A where *n*_*i*_ represents the number of supercoils in the *i*^*th*^ branch. Note that this solution satisfies the unknotted substrate equation for any choice of integers *n*_1_, *n*_2_, *n*_3_ as shown in Fig. 3B. If topoisomerase acts on a pair of loops emanating from a 3-branched structure, then the product will likely be a twist knot, a knot where supercoils are trapped by a clasp as shown in Fig. 3C.

Per proof in mathematical methods, if topoisomerase acts on each pair of loops emanating from a protein complex binding three DNA segments producing twist knots with less than 1000 crossings, then the only biologically plausible model for the DNA bound within the protein complex is a 3-branched supercoiled structure as shown in Fig. 3A. If knot types other than twist knots are produced via a single round of topoisomerase action, then the configuration will be more complicated. Potential models in this case can be determined computationally (Darcy et al., 2006). Thus we will focus on twist knot products.

**Fig. 3.**
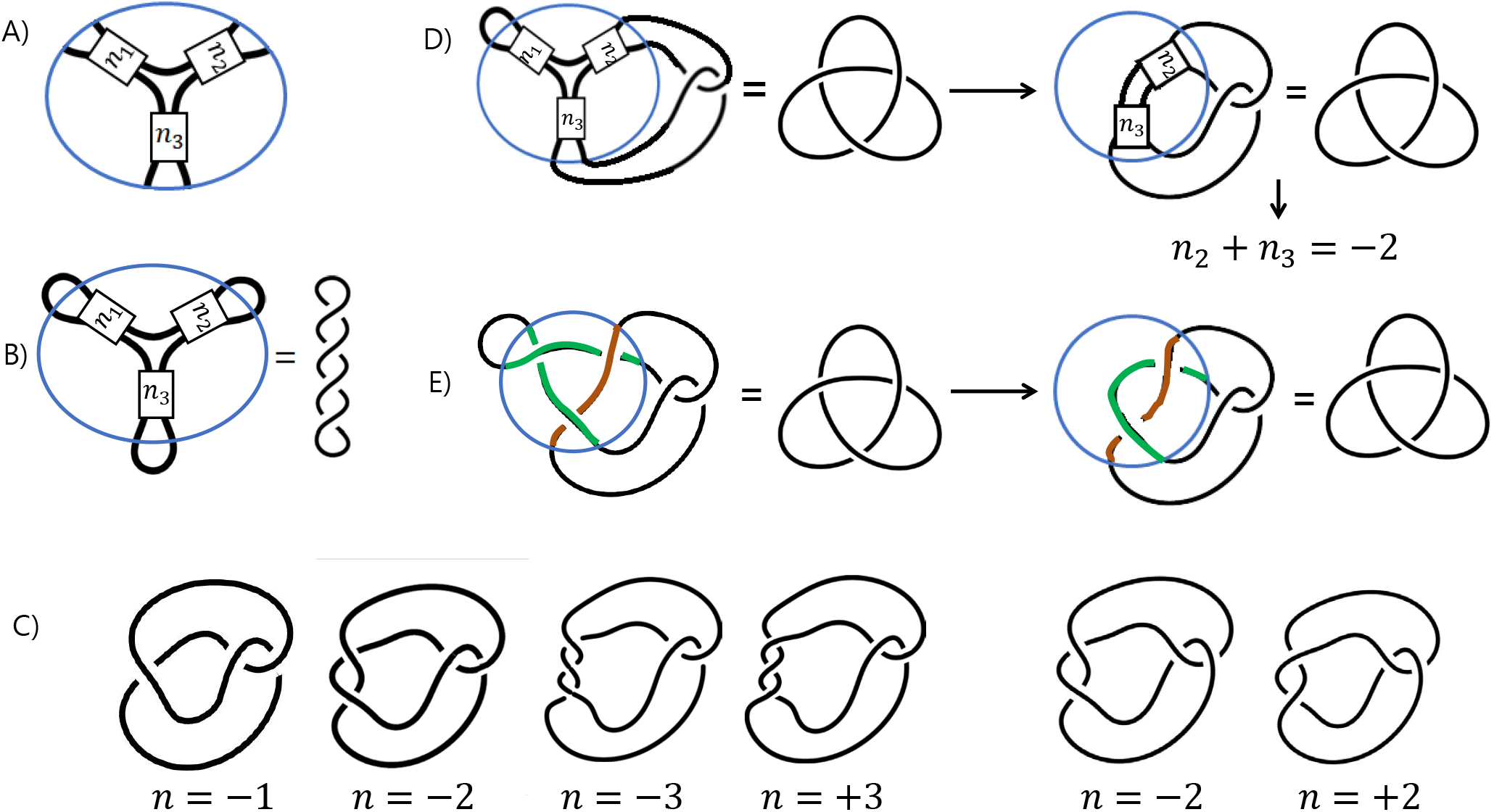
(A) A protein complex binding three DNA segments which form a 3-branched supercoiled DNA where *n*_*i*_ represents the number of supercoils in the *i*^th^ branch. The integer *n*_*i*_ is negative if intertwining is right-handed (and thus represents negative supercoils). If the intertwining is left-handed, then *n*_*i*_ is positive (and thus represents positive supercoils). (B) Unknotted subtrate equation assuming protein complex binds a 3-brnched supercoiled structure. (C) A twist knot is formed by trapping crossings via a clasp. If the clasp is left-handed and *n* = −1, then the knot diagram can be simplified to contain no crossing and thus we have the unknot. If *n* = −2, then the diagram can be simplified to the right-handed 3-crossing knot. If *n* = −3, then the diagram simplifies to the 4-crossing knot, while if *n* = 3, we obtain a 5-crossing twist knot. If the clasp is right-handed, then if *m* = −2, we obtain the 4-crossing knot, while if *m* = 2, then the diagram simplifies to the left-handed 3-crossing knot. (D) A 3-string tangle equation can be transformed into a 2-string tangle equation by pushing one of the outside loops into the tangle ball. In the case shown, we can then remove the supercoils in the branch containing *n*_1_ supercoils, leaving *n*_2_ + *n*_3_ supercoils. (E) An example of a solution to the 3-string tangle equation in (D) where *n*_1_ = *n*_2_ = *n*_3_ = −1. Note that since *n*_2_ + *n*_3_ = −2, the brown DNA segment crosses the green DNA segment two times.

#### Example 5

Consider the tangle equation in Fig. 3D where the product is the right-handed 3-crossing knot. Since topoisomerase acts on the loops on the right, the knotted product has *n*_2_ + *n*_3_ supercoil crossings between the clasp, while the *n*_1_ crossings in the loop on the left can be "mathematically removed" – i.e., removing these *n*_1_ crossings does not change the knot type of the product. After the protein complex is removed, the knot diagram can be simplified to a diagram with only three crossings. One solution to the tangle equation is shown in Fig. 3D. In this figure, we chose *n*_1_ = *n*_2_ = *n*_3_ = *−*1. We could have chosen any value for *n*_1_ and any pair of values for *n*_2_ and *n*_3_ satisfying *n*_2_ + *n*_3_ = *−*2. We know that *n*_2_ + *n*_3_ = *−*2 from Eqns 1 and 3 in Table 1. By removing the branch on the right, we have converted the 3-string tangle equation (Fig. 3D left) into a 2-string tangle equation (Fig. 3D right). Thus we can use Table 1 Eqn 3 to determine that *n*_2_ + *n*_3_ must represent two negative supercoils and thus *n*_2_ + *n*_3_ = *−*2. This also tells us how two of the three DNA segments interact. In Fig. 3E, the brown DNA segment must cross the green DNA segments two times. This will be true for all solutions since *n*_2_ + *n*_3_ = *−*2.

If we have three tangle equations, one for each pair of loops, that will give us three independent linear equations involving *n*_1_, *n*_2_, *n*_3_ for which there will be a unique solution. However, we don’t actually know which system of tangle equations best model difference topology experiments involving a topoisomerase. In particular, perhaps topoisomerase action results in the left-handed clasp (Fig. 2C middle) instead of the right-handed clasp (Fig. 2C right). In that case, equation 2 in Table 1 implies *n*_2_ + *n*_3_ = *−*1. Suppose the right-handed 3-crossing knot shown in the tangle equation Fig. 3D is the only product of a difference topology experiment involving topoisomerase. Suppose further that topoisomerase acted on every pair of loops, producing only right-handed 3-crossing knots. If we don’t assume which clasp, we obtain the following system of equations: (*n*_1_ + *n*_2_ = *−*2 or *n*_1_ + *n*_2_ = *−*1) and (*n*_1_ + *n*_3_ = *−*2 or *n*_1_ + *n*_3_ = *−*1) and (*n*_2_ + *n*_3_ = *−*2 or *n*_2_ + *n*_3_ = *−*1). We are only interested in integer solutions since in a projection, we cannot have a fractional crossing. The integer solutions to this system of equations are (*n*_1_*, n*_2_*, n*_3_) = (*−*1*, −*1*, −*1), (*−*1*, −*1, 0), (*−*1, 0*, −*1), (0*, −*1*, −*1). All these solutions could correspond to different projections of the same 3-dimensional model.

### Multiple DNA segments bound by protein via difference topology using topoisomerase

If a protein complex *P* binds four DNA segments, then a branched supercoiled structure might look like that in Fig. 4A. This configuration will be referred to as *0-standard*. However it is possible that the structure is more complicated as shown in Fig. 4B. This configuration will be referred to as *R-standard*. An R-standard tangle can be created from a 0-standard tangle by intertwining the branches containing *n*_1_, *n*_3_, *n*_4_ supercoils. For example, the tangle in Fig. 4B was created from the 0-standard tangle by first twining the branches containing *n*_1_ and *n*_4_ supercoils once, followed by twining the branches containing *n*_3_ and *n*_4_ supercoils twice. As proved in mathematical method section, if topoisomerase acts on each pair of loops emanating from a protein complex binding four DNA segments producing twist knots with less than 1000 crossings, then the DNA bound within the protein complex must be R-standard. Since 0-standard is more biologically relevant, we will focus on 0-standard solutions.

**Fig. 4.**
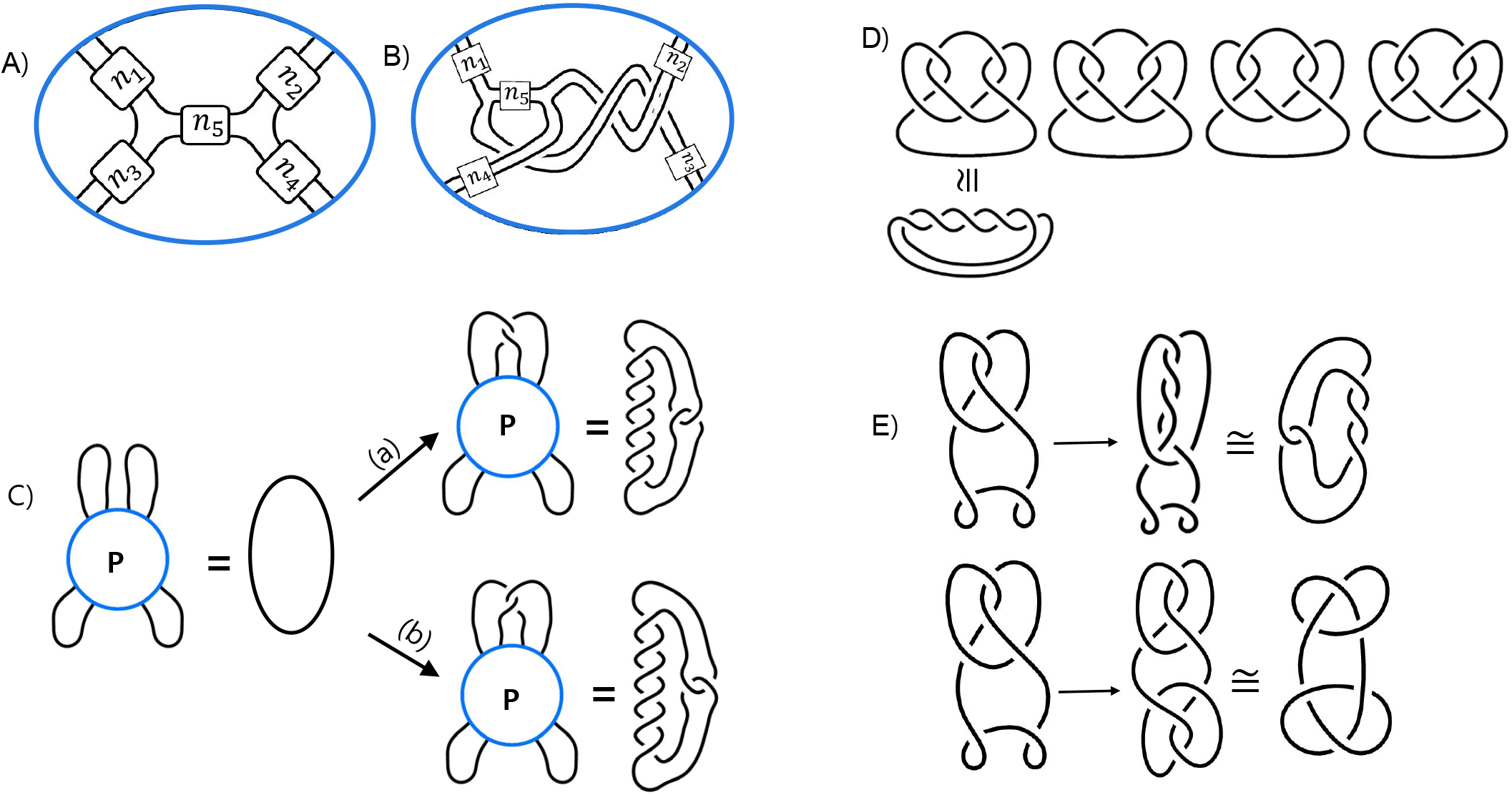
(A) A possible branched supercoiled DNA structure where the protein-complex has bound four DNA segments. This conformation with be referred to as 0-standard. The number of half twists in each branch is denoted by *n*_*i*_ where the half twists are left handed if *n*_*i*_ > 0 (representing positive supercoils) and right-handed if *n*_*i*_ < 0 (representing negative supercoils). (B) A more complicated branched supercoiled DNA structure. This type of conformation with be referred to as R-standard. (C) 4-string tangle equations modeling difference topology experiments with type II topoisomerase. The tangle *P* represents the 4-string tangle model of protein-bound DNA. Topoisomerase action will result in either (a) a left-handed clasp or (b) a right-handed clasp. (D). If topoisomerase acts on three consecutive loops, other knot types can be obtained. From left to right: If two left-handed clasps are created, then the result is a 5-crossing torus knot. If one left-handed clasp and one right-handed clasp are formed, then the product is a six crossing knot (middle two diagrams). If both clasps are right-handed, then the product is a seven crossing knot. (E) If topoisomerase acts twice on the same pair of loops, the result is a 5-crossing twist knot (top row). If topoisomerase acts on two different pairs of loops, then the result is the 6-crossing granny knot (bottom row).

We will use 4-string tangle analysis to analyze experiments involving condensins. Note that condensins bind multiple DNA segments. However, one can use 4-string tangle analysis to determine how any quadruple of strands interact. Moreover, often one can partition an *n*-string tangle model into smaller 4-string tangle models.

### Condensins and tangle equations involving multiple DNA segments

*Condensins* are large protein complexes that play a major role in chromosome assembly and segregation during meiosis and mitosis. Condensins interact with multiple sites of DNA to construct the structure of chromosome (Petrushenko et al., 2006; Uhlmann, 2016). *13S condensin* is a eukaryotic condensin from Xenopus. To understand how 13S condensin interacts with DNA, Kimura et al. used the difference topology experimental technique with type II topoisomerase (Kimura et al., 1999). Similar experiments were performed with *MukB*, the first discovered bacterial condensin in *Escherichia coli*, by Petrushenko et al. (Petrushenko et al., 2006). But they did not formally describe tangle equations modeling these reactions as the modeling is more complex for difference topology experiments and the mathematics for solving such equations did not exist until now.

In both the 13S condensin and MukB difference topology experiments, a nicked unknotted DNA substrate was used. This means that the DNA was not supercoiled. In the presence of type II topoisomerase and 13S condensin, the main product topology of DNA was the right-handed 3-crossing knot. In addition, they found a few four, five, and six-crossing knots. But as measured by gel electrophoresis, the 3-crossing knots were far more abundant than the higher crossing knots. For MukB the right-handed 3-crossing knot was also the main knot type produced in the difference topology experiment. But in this case, they also found a fair amount of four-crossing knots and some five-crossing knots.

To determine the handedness and knot type of the higher crossing products, some of the knots were identified via EM. Note EM does not quantify the amount of each knot type, but can be used to determine which knot type is more common among knotted products having the same crossing number if a sufficient number of knots are identified. For 13S condensin, of the 3-crossing knots identified, 130 were right-handed while only 8 were left-handed (Kimura et al., 1999). For MukB, 37 right-handed 3-crossing knots were identified while only one left-handed 3-crossing knots was observed (Petrushenko et al., 2006). Thus since most knots were determined to be 3-crossing knots via gel electrophoresis, and the vast majority of these knots were identified as right-handed via EM, we know that the main product of topoisomerase action on DNA bound by condensin are right-handed trefoil knots in both of these experiments.

## Results

Condensins bind DNA at multiple sites, so condensin-DNA complexes can be modeled by an *n*-string tangle, where *n* is a large number. We will focus on *n* = 4 since an *n*-string tangle model can sometimes be partitioned into 4-string tangle models. There are various hypothetical models of condensin-DNA complexes (Eeftens et al., 2017; Ganji et al., 2018; Hirano, 2012; Hayama et al., 2013; Kumar et al., 2017; Kimura et al., 1999; Losada and Hirano, 2005; Petrushenko et al., 2010; 2006; Rybenkov et al., 2014). Many of these models show a 4-string tangle model (for example Fig. 7 in (Petrushenko et al., 2006) as this model can be easily be extended to *n*-string tangle models.

### An n-string tangle analysis of condensin-DNA complexes

We will look at two cases. For the first case, we will assume the product of one round of topoisomerase action produces only right-handed trefoil knots. This is likely the case in the 13S condensin experiments as this was by far the major product based on gel electrophoresis and EM. For the second case, we will also assume that the right-handed trefoil knot is also the major product, but that products also included a fair number of four-crossing knots per the MukB experiments. Other products will be explained below in the minor products subsection. We focus on 0-standard solutions as other solutions are more complicated and hence unlikely to be biologically relevant.

#### 13C condensin

Suppose that 13S condensin binds DNA in a 0-standard configuration (Fig. 4A), and topoisomerase action on each pair of loops results in the right-handed trefoil knot. In this case, at least three of four branches (corresponding to *n*_1_, *n*_2_, *n*_3_, *n*_4_) contain one positive supercoil with the remaining branch containing one positive or no supercoils while the middle branch (corresponding to *n*_5_) will have one negative or no supercoil. These solutions can all be seen in the 3-dimensional model shown in Movie 3.

Note this solution is consistent with the models proposed in (Petrushenko et al., 2006), but we have additional information. If the solution is non-planar, then the twisting between branches should be right-handed since *n*_5_ = 0, −1.

#### MukB

Suppose that MukB binds DNA in a 0-standard configuration (Fig. 4 A), and topoisomerase action on each pair of loops results mostly in the right-handed trefoil knot as well as a fair amount of 4-crossing knots. If we restrict to the case where we assume at most 1*/*3 of the product consists of 4-crossing knots and the remaining products are right-handed trefoil knots, then there are 84 solutions. Several 2D solutions can be visualized via a single 3-dimensional solution, but we would still have a number of possible 3D solutions if we just considered mathematics. However, from a biological perspective, simpler solutions are likely to be more relevant. The simplest solution would be a modification of the solution we found for 13S condensin.

Consider the conformation shown in Movie 3. For topoisomerase action to result in the 3-crossing knot, the crossing change must result in the left-handed clasp (top pathway in Fig. 4C). In (Petrushenko et al., 2006), it was hypothesized that a major conformation change was responsible for the 4-crossing knotted product, but a conformation change is not needed. If topoisomerase action results in a right-handed clasp (bottom pathway in Fig. 4C), then a 4-crossing knot will be produced. The topoisomerase used in (Kimura et al., 1999; Petrushenko et al., 2006) does not have a chirality bias. Thus the fact that 3-crossing knots are more abundant than 4-crossing knots means that the loops must extrude from the protein complex in a manner such that a left-handed clasp is more likely to form than a right-handed clasp. This is particularly true for 13S condensin where the 3-crossing products were 20-fold more common than the 4-crossing products as detected via gel electrophoresis. Since 4-crossing products are more common for MukB (though still less than trefoils), this means that while both 3-dimensional models may project to the same configuration, there is a difference between their 3-dimensional configurations. Computational looping studies could be used to investigate 3-dimensional models more thoroughly (Vetcher et al., 2006).

#### Minor products

Note the model in Movie 3 can also be used to explain the minor products in the MukB and 13S condensin experiments. In both these experiments, a small number of five and six crossing knots were observed via gel electrophoresis (Kimura et al., 1999; Petrushenko et al., 2006). A few of the five and six crossing knots were identified via EM in (Kimura et al., 1999). Nine of the identified 5-crossing knots were right-handed 5-crossing torus knots (Fig 4D left). Four 5-crossing twist knots were also identified, one with the handedness shown in Fig 4E top and three with the opposite handedness. All five of the six crossing knots that were identified were right-handed granny knots (Fig 4E bottom). Since only a few five and six crossing knots were identified, we do not have a statistically significant sample size to infer which of these minor products were more common.

It is mathematically impossible for topoisomerase to create a 5-crossing torus knots or 6-crossing granny knots by acting only once on an unknotted substrate (Darcy and Sumners, 1997; Torisu, 1998). The 5-crossing torus knot will result if topoisomerase action occurs on two pairs of three adjacent DNA loops as in Fig. 4D left. If topoisomerase acts on two pairs of loops involving four DNA loops as in Fig. 4E bottom, the result is the 6-crossing granny knot. While the 5-crossing twist knot could be the result of a single round of topoisomerase action, it could also be the result of a second round of type II topoisomerase action on the same pair of loops used in the first round, see Fig. 4E top.

Note that for these knots, topoisomerase action results in left-handed clasps. If topoisomerase action also created right-handed clasps, then a 6-crossing knot, and a 7-crossing knot can be obtained as shown in Fig. 4D. However, those knots are not observed in the difference topology experiments of condensin-DNA complexes(Kimura et al., 1999; Petrushenko et al., 2006) supporting the preference for the formation of the left-handed clasp (Fig 4C(a)) over the right-handed clasp (Fig. 4C(B)) via topoisomerase action.

Some knots could also be the result of topoisomerase action on naked DNA. While knots were not detected in the control reaction where topoisomerase acted on naked DNA, gel electrophoresis may not detect all knots. There could be some bands containing knots that are not visible via Southern blotting. Thus the small number of left-handed trefoils and five-crossing twist knots could result from topoisomerase action on naked DNA.

### Discussion

While tangle equations related to difference topology experiments with Cre have been well studied (Pathania et al., 2002; 2003; Yin et al., 2005; Harshey and Jayaram, 2006; Darcy et al., 2006; Yin et al., 2007; Darcy et al., 2009), difference topology experiments using type II topoisomerase have not yet been actively studied. We analyzed the difference topology experiments of condensin-DNA complexes with type II topoisomerase (Kimura et al., 1999; Petrushenko et al., 2006) by using the tangle model, and conclude that the conformation shown in Movie 3 is the most biologically relevant tangle model for condensin-DNA complexes. This conclusion also supports the working model suggested by Petrushenko et al. (Petrushenko et al., 2006).

Note that in both the model in (Petrushenko et al., 2006) and our model based on the data in (Petrushenko et al., 2006), condensin binds to positive supercoils. If condensin bound only negative supercoils, then topoisomerase would not have produced right-handed 3-crossing knots. When a protein-complex binds DNA, the DNA may partially unwind where the protein-complex binds DNA, resulting in local undertwisting of this protein-bound DNA. DNA in the cell is generally negatively supercoiled. This negative supercoiling allows for proteins to more easily untwist the two strands of double-stranded DNA. When this occurs, the negative supercoiling is converted to underwound DNA. Thus the amount of negative supercoiling decreases. This can occur via a decrease in the number of supercoils or in the creation of a positive supercoil. When condensin binds a DNA loop, the local untwisting of DNA can result in a positive supercoil in that loop. Condensin bound to that positive supercoil would trap that supercoil and thus topoisomerase action results in a right-handed 3-crossing knot.

If the change in DNA twist caused by the binding of condensin is offset by the creation of a positive supercoil bound by condensin, then there would be no change in the overall supercoiling of the DNA. But the supercoiling bound by protein need not match the change in twist caused by protein binding. For both MukB (Petrushenko et al., 2006) and 13S condensin (Kimura et al., 1999), a net change in supercoiling was observed. In the case of MukB, the net change was negative, while for 13S condensin, the net change was positive. Recall that the middle branch in the 0-standard configuration will contain either 0 or 1 negative supercoil (represented by *n*_5_ in Fig 4A). If the middle branch contains a negative supercoil, this could explain the net negative change in supercoiling for MukB. The tangle equations give us possible 2-dimensional models for protein-bound DNA. Per the movies, this is one reason that we often obtain more than one solution to systems of tangle equations modeling difference topology experiments involving a topoisomerase: the actual 3-dimensional model can project to more than one 2-dimensional tangle solution. In particular the 3-dimensional conformation of DNA is unlikely to have exactly 0 or exactly 1 negative supercoil in the middle branch represented by *n*_5_. Thus in three dimensions, *n*_5_ would be better represented by a fraction between 0 and −1. We hypothesis that for MukB, the 3-dimensional conformation of DNA bound in the MukB-DNA complex contains a middle branch with a fractional negative supercoil closer to one, while for 13S condensin this fractional negative supercoil would be closer to zero. This would explain both the difference in knotted products as well as the net change in supercoiling (negative for MukB and positive for 13S condensin).

The main advantage of difference topology experiments using a topoisomerase is that one can determine the feasibility of this experimental technique applied to a particular protein complex without a significant investment in time. If one uses a site-specific recombinase, one must create substrates with recombinase binding sites correctly placed on every pair of loops. If one uses a topoisomerase, one can simply add topoisomerase to a test tube containing the protein complex under study using reaction conditions where this protein complex stably binds DNA. The control reaction also must be performed where topoisomerase acts on naked DNA under identical reaction conditions except for the omission of the protein complex under study. If there is a difference in the knot types of the products as determined via gel electrophoresis, then difference topology can give insight into the conformation of DNA bound by protein. Moreover, depending on difference topology experimental results using a topoisomerase, one can determine whether one is likely to obtain sufficiently better information using a recombinase instead of a topoisomerase. One can also use the difference topology results using a topoisomerase to predict the results if one were to use a recombinase instead (Darcy and Price, to appear).

Of course, experimental procedures are rarely simple. One may need to play around with reaction conditions. In particular, one might need to use a singly nicked DNA substrate as was used in the condensin experiments. If supercoiled DNA is used, then topoisomerase may trap supercoils not bound by protein. One can use the simplest products to determine the minimal complexity of DNA bound by protein, but results involving multiple different types of DNA knots would be much harder to analyze. Using a single nicked DNA may reduce the number of knot types resulting from topoisomerase action. However, the use of a nicked DNA substrate can affect the DNA conformation bound by DNA. Recall that difference topology was used to determine that Mu transposase binds to five DNA crossings per the tangle model in Fig 1A. In this case a supercoiled DNA substrate was used. When difference topology with Cre recombinase was applied to Mu transposase binding to nicked DNA, Mu transposase only bound four DNA crossings instead of the five DNA crossings. (Yin et al., 2005).

While analyzing difference topology experiments involving a topoisomerase is not as straight-forward as analyzing difference topology experiments involving a site-specific recombinase, the results can still be very useful for modeling the 3-dimensional conformation of DNA bound by protein as we have illustrated with our analysis of 13S condensin and MukB. When tangle equations have been used to model a difference topology experiment using a site-specific recombinase, only one biological relevant 2-dimensional tangle solution has been found (Pathania et al., 2002; 2003; Yin et al., 2005; Harshey and Jayaram, 2006; Yin et al., 2007). The ambiguity in the tangle model for difference topology experiments involving a topoisomerase means we do not expect to find only one biological relevant 2-dimensional tangle solution. But this ambiguity led us to consider 3-dimensional models consistent with the 2-dimensional tangle solutions. So while difference topology experiments involving a recombinase are less ambiguous and easier to analyze, we obtain better 3-dimensional analysis when a topoisomerase is used instead.

## MATERIALS AND METHODS

### Mathematical Methods

Most 2-string tangle equations modeling topoisomerase action can be solved using the TopoICE-X (Darcy et al., 2008) in KnotPlot (Scharein, 1998) including all equations where the knots involved have fewer than 8 crossings.

We use the following theorem to solve 3- and 4-string tangle equations:

#### Theorem 1.

*Suppose protein P binds DNA and topoisomerase acts on all pairs of loops producing only twist knots with fewer than 100,000 crossings. Then the only biologically relevant tangle model representing P is a 3-branched structure* (*Fig* 1 *A*) *if protein P binds to three DNA segments or P is R-standard if protein P binds to four DNA segments*.

*Proof*: Consider the tangle equation modeling topoisomerase acting on one pair of loops emanating from the protein *P*-DNA complex. We can convert this tangle equation into a 2-string tangle equation by pushing all loops except the pair of loops upon which topoisomerase acted into the tangle ball (see for example Fig 3D). We can solve the 2-string tangle equation using results in (Torisu, 1998; Darcy and Sumners, 1997) where the product of topoisomerase acting on unknotted DNA is a twist knot. We solved all such equations where the twist knot had fewer than 100,000 crossings by writing a simple C program (code available upon request). This allows us to use the results in (Darcy et al., 2009; Kim and Darcy, 2015) to prove that the only biologically relevant tangle model representing *P* is a 3-branched structure if *P* binds three DNA segments or *R*-standard if *P* binds to four DNA segments. □

## Acknowledgements

We thank Rob Scharein for drawing all figures and creating movies by using KnotPlot.

## Contribution

Conceptualization: I.D., S.K..; Methodology: I.D., S.K.; Software: I.D..; Formal analysis: I.D., S.K..; Writing: I.D., S.K.

---

**Movie 1:**
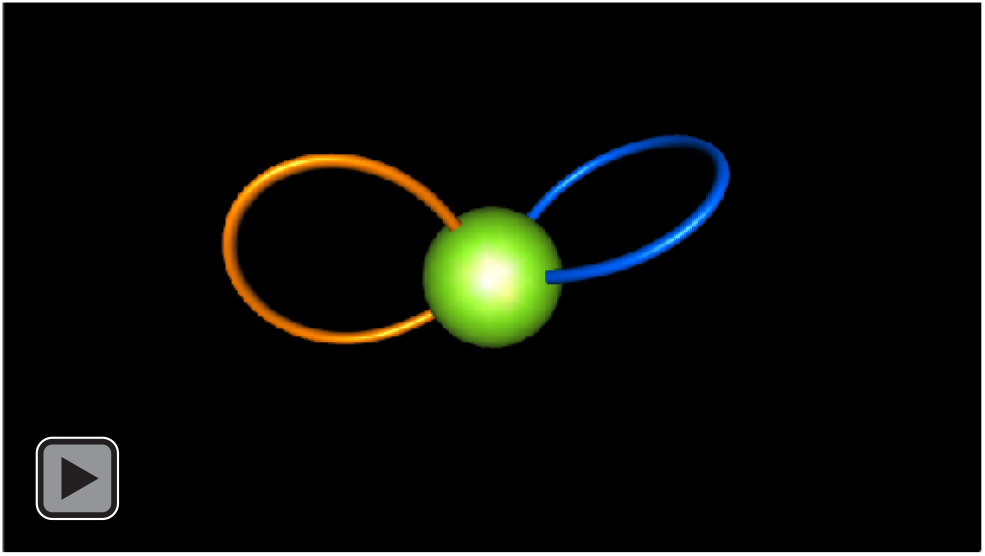
Exterior loops. Depending on how one projects a 3-dimensional protein-DNA complex, the two outside loops may or may not cross.

**Movie 2:**
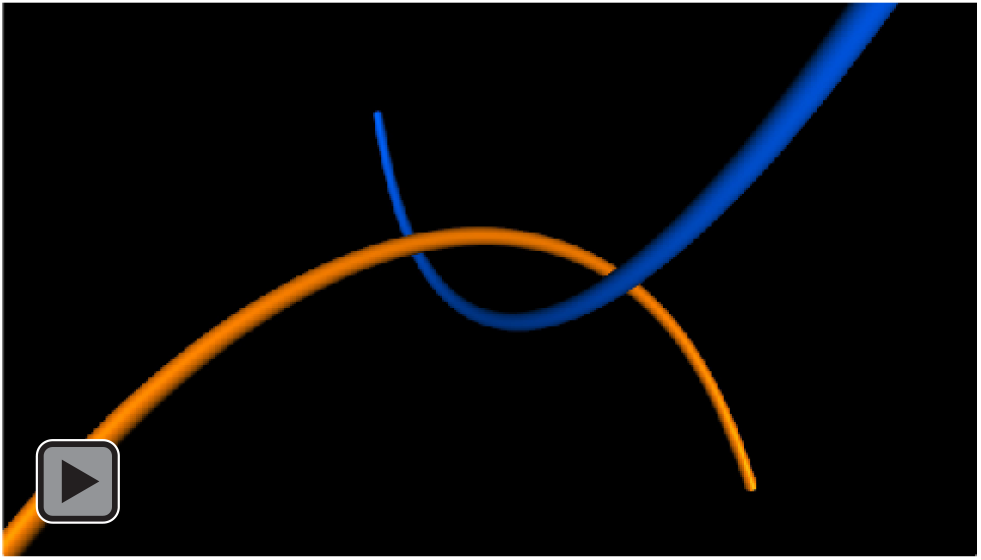
Left-handed vs right-handed clasp. The clasp first appears as a left-handed, but viewing the clasp from the other side, the clasp appears as a right-handed clasp. Thus whether or not a clasp is right-handed or left-handed is projection dependent. In this movie the two segments cross at a 90 degree angle. But if the two segments cross at a different angle, than one handedness will appear in a larger percentage of projections than the other handedness.

**Movie 3:**
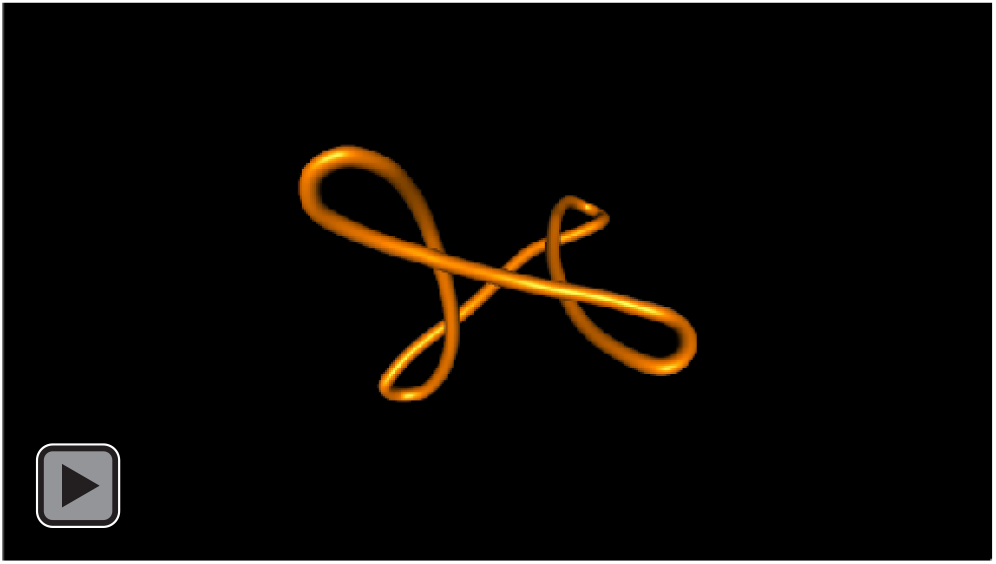
4-string tangle model for DNA bound by condensin. The 4-string tangle model contains one postive supercoils in each of the four branches and one negative supercoil in the middle branch. As we rotate this model, the negative supercoil in the middle is no longer visible and we briefly see only three positive supercoils. As we continue rotating four positive supercoils appear. We again briefly loose one positive supercoil until we return back to the projection where we can view all four positive supercoils and the one negative supercoil in the middle branch.

